# Data Resources and Analyses Fair Header Reference genome: A Trustworthy standard

**DOI:** 10.1101/2023.11.29.569306

**Authors:** Adam Wright, Mark D Wilkinson, Chris Mungall, Scott Cain, Stephen Richards, Paul Sternberg, Ellen Provin, Jonathan L Jacobs, Scott Geib, Daniela Raciti, Karen Yook, Lincoln Stein, David C Molik

## Abstract

The lack of interoperable data standards among reference genome data-sharing platforms inhibits cross-platform analysis while increasing the risk of data provenance loss. Here, we describe the FAIR-bioHeaders Reference genome (FHR), a metadata standard guided by the principles of Findability, Accessibility, Interoperability, and Reuse (FAIR) in addition to the principles of Transparency, Responsibility, User focus, Sustainability, and Technology (TRUST). The objective of FHR is to provide an extensive set of data serialisation methods and minimum data field requirements while still maintaining extensibility, flexibility, and expressivity in an increasingly decentralised genomic data ecosystem. The effort needed to implement FHR is low; FHR’s design philosophy ensures easy implementation while retaining the benefits gained from recording both machine and human-readable provenance.

## INTRODUCTION

### Importance of reference genomes

The large and ever-increasing number of well-characterised reference genomes has become a prerequisite for many essential analyses, including cross-species comparisons and population genomics studies of model and non-model systems (1). Unfortunately, there are no file header standards for the metadata describing these essential resources, and even basic fields, such as “species” and “strain”, are missing from the common reference genome data standards. Instead, such metadata must be kept separately from the files containing the reference genome data, raising the risk of gaps in data provenance and human copying errors and imposing a burden on computational biologists and developers of analytic software alike (2, 3). The FAIR-bioHeaders Reference genome (FHR) specification aims to provide a standard to maintain the provenance of reference genomes that is translatable across storage and analysis platforms.

### History of FASTA

The FASTA file format is widely used in genomic analysis as the repository of reference genome sequence information. FASTA was developed in 1985 (4) and is still widely used for reference genomes, but during that time the genomics ecosystem has changed dramatically; the first journal mention of a centralised DNA repository was in Elke Jordan and Christine Carrico’s Science Letter in 1982 (5). The FASTA format specification (originally the “Pearson format”) was created by William Pearson and David Lipman in 1985 (4), but has since been maintained by the National Center for Biotechnology Information (NCBI) at the U.S. National Institutes of Health (4).

It used to be that reference genome files resided in a small number of trusted repositories, such as GenBank, and were downloaded from there to the bioinformatician’s local computer system, it is increasingly the case that these files are modified, reannotated, and redistributed in a decentralised manner (6, 7, 8). This was not the consequence of any deliberative process. Instead, it occurred organically as a consequence of the shared ownership of the reference data and the collaborative nature of science (9). Decentralisation also occurs when multiple websites host genomic data instead of a single authority, and it is unlikely that this process will reverse (6). Decentralisation carries risks, not least of which is the loss of provenance metadata that may occur when the files are transferred among resources and when users download the genomes for local processing. Furthermore, loss of provenance reduces the ability of users to ensure that data are what they claim to be, potentially causing confusion, propagating errors in subsequent analysis, and increasing overall time and effort to reuse data (10).

### Deficiencies of FASTA for reference genomes

It is highly problematic that reference genome FASTA files contain no intrinsic information that describes the nature and provenance of their contents; all provenance information must come from external sources and be linked to the file name or checksum. However, both of these methods are prone to loss of information. Files can be easily renamed or overwritten, and when the name has changed, the link to provenance information can be difficult to recover. Checksums, an algorithmically unique representation of a file that can be compared for accuracy, are also brittle. Commonly performed file manipulations, such as introducing carriage returns when a file generated on a Linux system is opened in a Windows text editor, introduce alterations that do not affect the semantics of the file, but completely change the checksum. By relying on external information for the provenance of the file, bioinformaticians risk associating incorrect metadata with the genome file or even being unable to locate the metadata at all (11, 12).

Differences can arise when a reference genome is replicated across platforms or devices (e.g. renaming of files or contigs, removal of contigs that fail to meet some criteria such as minimum length, the removal and addition of metadata, etc.) leading to a gradual divergence of reference genome files and their metadata (i.e., the genome data and metadata divergence problem, divergence problems are described by Haslhofer 2010 (13)). Furthermore, discrepancies can arise when a genome assembly is updated with additional data, due to inexact version matching from multiple genome assembly versions and user updates across platforms. To address the discrepancies that arise from replication, what could be called a reference genome authority is typically implemented (e.g. https://www.ncbi.nlm.nih.gov/assembly, https://www.ncbi.nlm.nih.gov/genbank/, https://www.ddbj.nig.ac.jp/index-e.html), a central site that provides the authoritative version of a reference genome and its origin. While reference genome authorities are the ideal solution, in the current biological data environment, a reference genome authority is not always a practical solution. Two recent examples of reference genomes being published in multiple locations illustrate not only the need for genome hosting to be available, but also the necessity of decentralised assembly hosting and how file-level discrepancies can be introduced.

One example is the American Type Culture Collection (ATCC), a major biorepository and living culture collection that provides researchers with the physical strains and cell lines needed for their research. Historically, materials obtained from ATCC have been subjected to whole genome sequencing by researchers using those materials in their own research. The resulting genome assemblies produced by researchers are often submitted to the NCBI Assembly reference genome database in order to disseminate and share the data (14). The NCBI Assembly reference database, in this case, may be thought of as the reference genome authority. However, gaps in genomics data quality, data provenance and the traceability of materials used by researchers have contributed significantly to the scientific reproducibility crisis (reviewed in Hirsch 2019: (15)). In response to issues of provenance and authenticity, ATCC launched the https://genomes.atcc.org/ to establish its own quality control and provenance standards associated with genome references that represent authentic ATCC materials (16). ATCC has thus far produced over 4,000 high-quality or closed reference genomes for microbes within the ATCC collection, all under an ISO 9000 controlled quality assurance framework. This presents some dilemmas, however, as the NCBI Assembly database includes (for example) genome references for bacterial strains that have serious gaps in metadata or include substantial errors in their genome assembly when compared to the ATCC Genome Portal reference (17). Reducing discrepancies between genome references for the “same” organism can be aided by improving our ability to include crucial metadata about the origins of and means by which each genome reference is created in-line with the sequence data itself.

Another example of discrepencies that can arise from gaps in provenance can be found in molecular data portals and genome browsers. Several organism-focused genome data portals, such as AgBase (18), FlyBase (19), SoyBase (20), wFleaBase (21), WormBase (22), VectorBase (23), Ensembl (24), and others (25), publish annotations that are not found in the NCBI Assembly database. In some cases, these annotations and associated genomes cannot be submitted due to data ownership conflicts. These genome browsers and data repositories are often associated with a larger consortium that is working to answer questions of interest to the relevant scientific communities. Examples of such consortiums are the i5k (26, 27) Workspace (28), a collaborative effort to annotate arthropod genomes, and the Alliance of Genome Resources (The Alliance) (29) a centralised resource Model Organism resource. However, the data that these communities require have specific requirements, which can lead to the data portals and genome browsers becoming the primary source of their scientific communities’ reference genomes. Regardless of the resources that currently host the genome, there is no link to the source of the file, including its metadata, and associated publications. Since decentralisation is currently ongoing, it is unreasonable to imagine a world in which reference genome assemblies are not shared across platforms.

In this paper, we present the FHR FASTA header specification, which has been developed to address the genome data and metadata divergence problem. A key benefit of FHR is that it minimises the technical impact of adding provenance metadata to FASTA reference genome files by utilising legacy features of the file format instead of adding completely new ones. FHR is designed to enable FAIR and TRUST principles(30, 31), and to reduce the risk of data loss by ensuring that the provenance metadata is tied to the reference genome.

## METHODS

FHR version one is publicly available on GitHub within the organisation FAIR-bioHeaders. There are two relevant repositories: the specification and FHR-related tools (i.e. the FHR conversion and validation toolkit). The specification is codified within JSON following the JSON Schema specification. The related tools are written in Python and use this JSON schema to validate FHR-specified files. To ensure that the software is as maintainable as possible, we chose to use a minimal set of well-established dependencies. The Python libraries on which FHR converter tools depend include: YAML, JSON, Microdata v0.8.0, re (REGEX), hashlib, and JSONSchema v4.17.3 libraries. The tool requires Python 3.6 or higher and can be installed using the Python package setuptools version 42 or higher. These dependencies are all that are required to validate and convert FHR files. The goal of using so few dependencies is to reduce the potential for issues that arise when installing the tools and reduce the effort required to maintain the tools.

## RESULTS

The core constituent FHR fields were determined by considering the hypothetical, but unlikely, scenario of catastrophic loss of all copies of a reference genome. In this hypothetical scenario, all digital copies of the genome assembly have been lost, along with raw data. To reconstruct the genome, the following fields would be required: the location of the biological materials used to create the genome, the sequencing instruments used, and the assembly tools used to assemble and quality-check the genome. Therefore, FHR records the location of the biological materials, the sequencing instruments, and the assembly software tools. Furthermore, FHR records other information that would be useful in recovering such a genome: the metadata author of the FHR document, the assembler used, and other documentation either in the FHR instance or found in any scholarly articles associated with the genome. FHR also records funding and licencing information, related links, and the name and version of the genome. The intent of FHR is to strike a balance between only forcing users to provide the minimal information about a reference genome assembly but also be flexable enough to include other information. Therefore, the required fields are the absolute minimum to provide provenance of the data, and the optional fields focus on providing other useful information, and flexibility. It is recognized that more information on sample preparation and data processing could be added to the header but to keep the specification reasonably concise not all possible fields were added to the specification.

### Required fields

The FHR specification has nine required fields: `schema`, `schemaVersion`, `genome`, `taxon`, `version`, `assemblyAuthor`, `metadataAuthor`, `dateCreated`, and `checksum`. Please refer to (Table 1, and Fig. 1).

**Table 1.**
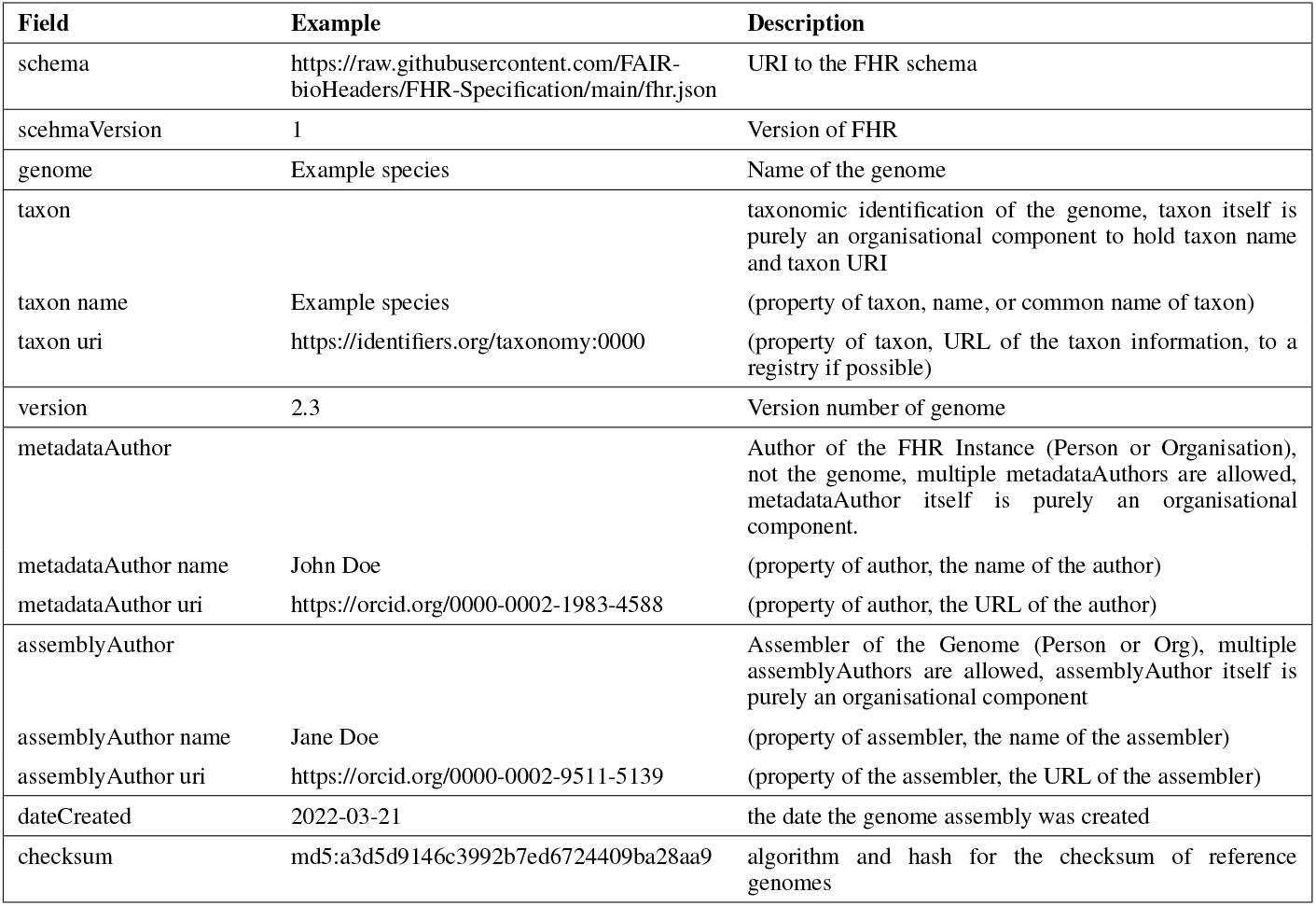
FHR Required Fields.

**Table 2.** FHR Optional Fields.

| Field | Example | Description |
| --- | --- | --- |
| assemblySoftware | Assembler.py | assembly software used in the creation of the genome |
| documentation | assembly of Example species | documentation about the genome |
| funding | Project number 1024512128 | grant line item, multiple funding lines are allowed |
| genomeSynonym | Other common names | Other names of the genome |
| identifier | eg:1024512128 | miscellaneous identifiers used to identify the genome |
| instrument | Awesome Sequencer IIe | physical tools and instruments used |
| reuseConditions | Creative Commons 4.0 | Specifying how the information in the file can be used |
| accessionID |  | physical ID of the genome assembly |
| accessionID name | BioSample:00000000 | (property of accessionID, the ID of the sample ) |
| accessionID uri | https://example.org/awesome_science/project-1024 | (property of accessionID, the URL of the physical sample) |
| voucherSpecimen | Located in Freezer 33, Drawer 137 at Cool Science Organisation Rm 1024 | a description of the physical sample |
| relatedLink | https://example.org/example_species/our_genome | related URLs to the genome, multiple relatedLinks are allowed |
| scholarlyArticle | https://doi.org/10.3390/insects12070626 | the genome of the scholarly article was published in |

**Figure 1.**
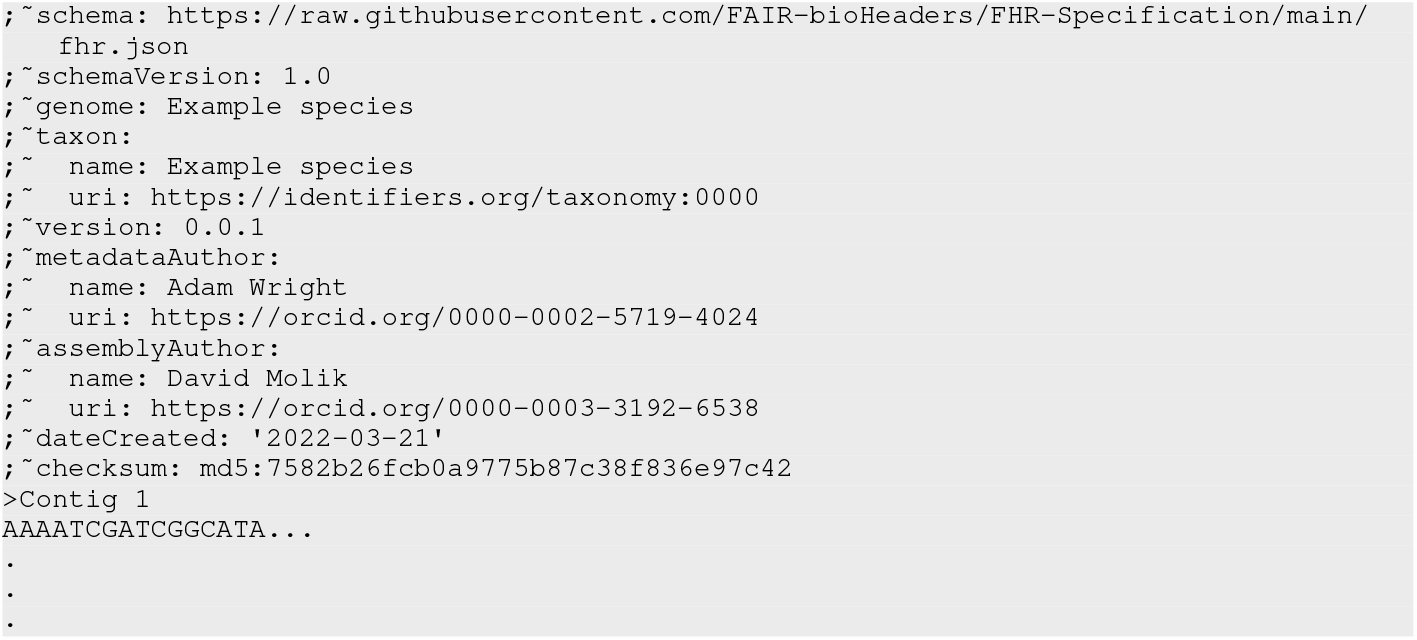
Minimal FHR FASTA header example with sequence

**Figure 2.**
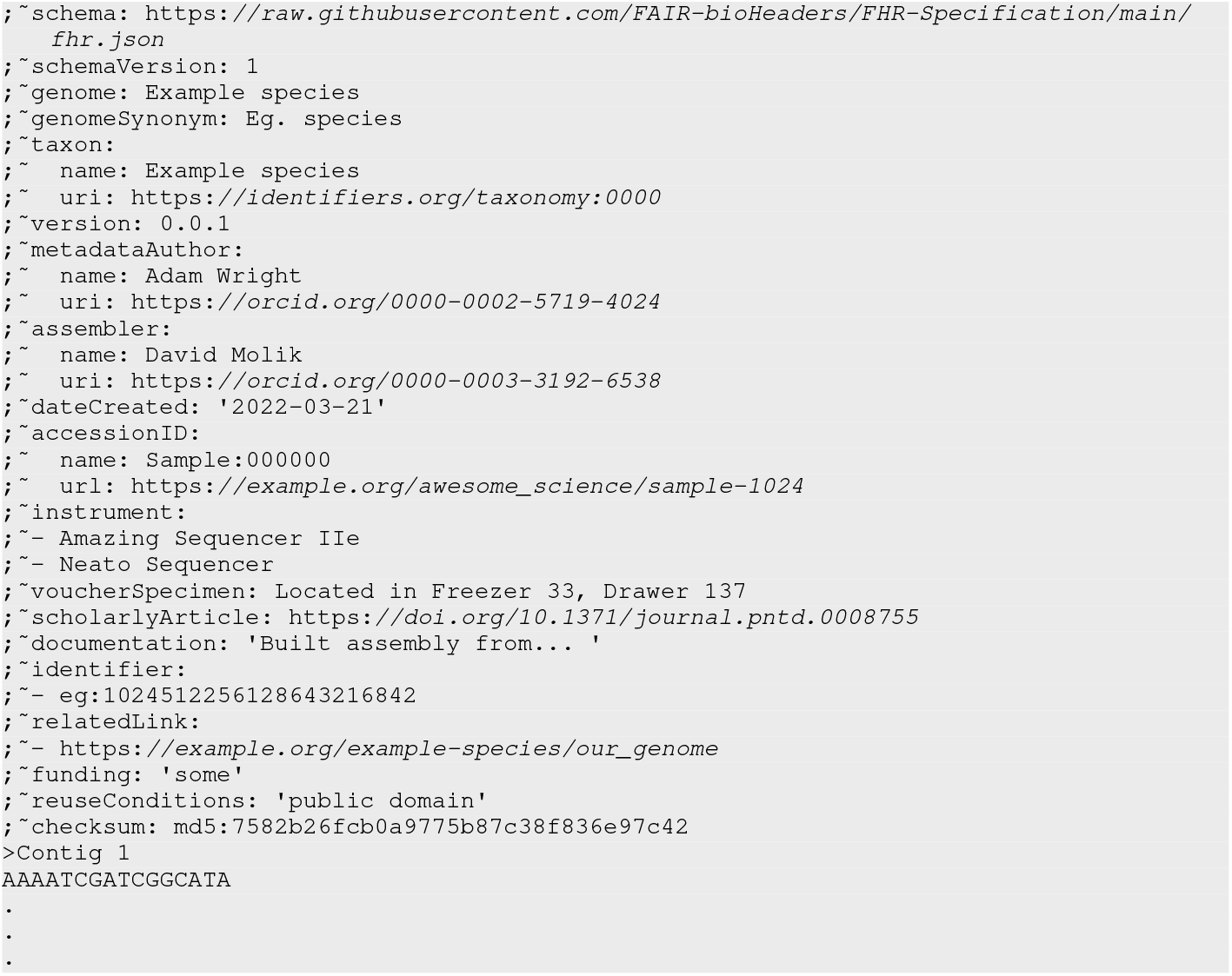
Expanded with optional fields FHR FASTA header example with sequence

The `schema` field indicates which JSON schema specification the metadata adheres to. In combination with the `schemaVersion` field, it allows users and software to know the exact format of the metadata to which the header conforms. The `genome` field is a string that is used to refer to the common name of the genome. The field contents are chosen by those generating the reference genome and can be a human-readable name, an alphanumeric ID, or a URI. The latter options are designed to simplify automated analysis.

`taxon` allows the user to specify the species in both a human-readable string as well as a link to identifiers.org to provide more information on the genome.

The `assemblyAuthor` field is a list of authors who participated in the generating of the genome; FHR supports multiple assembly authors. Typically, these authors would be those who contributed to the original paper(s) describing the sequencing and annotation of the genome, but this choice is left to those who generated the reference genome. In contrast, `metadataAuthor` identifies the person who authored the header metadata; FHR supports multiple metadata authors, as multiple individuals or organisations can contribute to the recording of its provenance. By providing both authorship fields, FHR supports situations where the creators of the FHR metadata are not the same as the creators of the assembly itself; this is useful when FHR metadata have been added after the fact by another group.

The `dateCreated` field specifies the creation date for the complete reference genome file. It works hand in hand with the `checksum` field, which provides a way of confirming that neither the data nor the metadata have changed since the file was created. We consider the checksum field to be one of the FHR format’s most useful features. In addition to being used to ensure that the file has not been corrupted, the checksum can also be recorded by the pipeline to keep track of the exact assembly used within an analysis run. Pipelines typically record the name of the assembly (e.g. HG19), but it is not uncommon for various *ad hoc* variants of the assembly to be circulated within the community, causing ambiguity. The checksum uniquely identifies the assembly and its metadata, thereby providing an identifier that can be used by analytic pipelines to unambiguously declare which assembly their results are based on.

Together, these nine fields allow researchers to identify the format of the metadata, identify the contents of the file, and keep track of who made the file and when, thereby enabling provenance-tracking.

### Optional fields

In addition to the nine required fields, FHR encourages the addition of up to 11 optional fields. These additional fields provide additional information on how the genome was assembled and distributed.

The `identifier`, `relatedLink`, and `scholarlyArticle` provide the user with links to external information about the genome that is not present in the FASTA file itself. The `identifier` field is used to associate compact URIs (“CURIEs”) with the reference genome. `identifier` is the main field from which a genome assembly can be mapped to a schema or database of an organisation. Both `relatedLink` and `scholarlyArticle` are conventional URLs. `relatedLink` is intended to associate the reference genome with its host site and mirrors, while `scholarlyArticle` is intended to link the genome to its marker paper. All these fields can take multiple values. FHR requires both URLs and identifiers to be publicly accessible and persistent. In particular, taxonomic information (i.e., `taxon`) such as species and isolate / cultivar must be identified using taxonomy URIs registered with identifiers.org in the `taxon uri` as well as the name of the taxon in the `taxon name`. When using CURIEs, FHR recommends that they be registered with identifiers.org so that the user can conveniently find the resource associated with the CURIE. As an alternative to identifiers.org, FHR allows Bioregistry (32) CURIEs to be used. If there is no suitable URI for use in an identifier field, the FHR specifications call for the use of the fields `relatedLink` and `documentation` described later.

The `assemblySoftware`, `instrument`, `accessionID`, and `voucherSpecimen` fields provide additional information on how the genome was generated. `instrument` is a multivalue field that refers to sequencing machines and DNA prep instrumentation used to generate the reference genome. `assembly software` is used to store the name and version of the assembly software used in the creation of the genome assembly, `accessionID` refers to the ID of the genome assembly, and `voucherSpecimen` is used to describe the location of the sequenced material.

Another optional field, `genomeSynonym`, can be used to add one or more common names to the reference genome to supplement the primary `genome` field. It is particularly useful when `genome` contains an unfriendly machine-readable identifier.

`documentation` can be used to provide additional information about the genome. It is free human-readable text that can be used to describe how the reference genome was made, its provenance, usage caveats, or any other information that users of this data should be aware of.

The final optional fields are `funding` and `reuseConditions`. `funding` allows the project funders to be acknowledged and can be multi-valued. The `reuseConditions` field can be used to specify the licencing terms under which genomic data can be used, if any. Although these fields are all optional, providing them helps increase the FAIRness and TRUSTworthiness of the reference genome. They were chosen to give the authors of the genome multiple complementary ways to document the contents of the file. For example, the `genome` ID field would typically be used to obtain information about the genome from external sources. However, in situations in which the genome’s ID is not sufficient on its own, the authors can add specific relevant information using the `assembly identifier`, `scholarlyArticle`, and `relatedLinks` fields.

### Full specification

The full formal specification of FHR is located in our GitHub organisation FAIR-BioHeaders in the repo FAIR-BioHeaders/FHR-Specification. It is versioned using GIT tags. Each release has a full change log, readily accessible through GitHub. Full documentation and examples accompany the FHR-Specification. FHR is specified using a JSON Schema file located at FAIR-BioHeaders/FHR-Specification/fhr.json. The FHR header’s `schema` field points to this file and is used by FHR software tools to verify the formatting and validity of the FHR header. In addition to the JSON schema, FHR has a human-readable specification and examples for FHR. The human-readable specification is located at FAIR-BioHeaders/FHR-Specification and includes both documentation and practical examples.

#### FASTA header

The recommended method for attaching FHR metadata to a FASTA file is to incorporate it into the FASTA file’s header. The FASTA format consists of a series of one or more nucleotide or protein sequence records, each preceded by a one-line header prefixed with ‘>’. The current schema for FASTA provided by NCBI defines the header line specification as consisting of a structured string that includes the Sequence Identifier (SeqID) followed by optional components such as the Resource ID, the accession number and the name of the Sequence (33). There is no formalized way to add additional information to the sequence-level header line (i.e. various resources utilize the sequence header differently).

FHR exploits a legacy feature of FASTA, the FASTA comment, which was included in the original specification created by Lipman & Pearson (1985). The comment is an optional multiline header that appears at the top of the FASTA file, each line of which is preceded by a semicolon (see Table 1). The FHR FASTA header consists of the metadata formatted according to the YAML specification, described below, and each line is prefixed by “; ∼“. This technique provides a format that is both easy to read and easily parsed computationally (34). Furthermore, this format fits into recommendations presented in Batista’s paper on “Machine actionable metadata models” (35).

The original FASTP tool, which introduced the file format of the same name and was published in 1985, supported FASTA comments (4, 36), and legacy tools that load and parse FASTA files are supposed to ignore these lines. Unfortunately, this is not always the case. Although some modern FASTA-consuming tools recognise and ignore semicolon-based FASTA comments, most do not. Fortunately, it is trivially easy to strip comments out of a FASTA file by removing lines that begin with semicolons. Users of FHR-enabled FASTA files may need to add this preprocessing step to their nucleic acid analysis pipelines before passing the file to downstream tools.

### Other serialisation methods

Although the FASTA header format is the preferred implementation of FHR, presenting and transferring FHR header may necessitate other FHR format serialisations. When not embedded directly in the FASTA header, FHR data can be associated with a reference genome by pairing it with a supplementary file that shares the same folder and base name as the FASTA file. This avoids issues with FASTA-consuming software tools that cannot parse the semicolon-delimited comments used by the FASTA FHR header. FHR supplementary files can be represented using several different file formats, JSON, YAML, and HTML, each specialised for a different use case. JavaScript Object Notation (JSON) is widely supported across multiple programming languages, and YAML Ain’t Markup Language (YAML) provides a format that is both machine- and human-readable, while Microdata embedded HTML is used to expose metadata to web search engines. We provide a conversion tool to inter-convert these formats.

We recommend naming the FHR file using the template `<genome file name>.fhr.<yaml|json|html>`. It should be located in the same filesystem directory as the FASTA file. In the case of FASTA files that are referenced using a URL, the FHR file should share the same URL path as its corresponding FASTA file. Following a standard naming system reduces the chances that the FASTA metadata becomes decoupled from the data or associated with the wrong data.

#### Checksum generation method

The checksum is computed based on the contents of the file, minus the checksum, which is injected into the header after being generated. In this way, the checksum can uniquely identify the reference genome sequences, as well as the header in the downstream analysis. For example, for a BAM file generated from an FHR-compliant reference genome, the reference genome can be uniquely identified through its FHR header’s checksum, facilitating accurate provenance tracking throughout the analysis pipeline. As all serialisations of FHR are formatted text files, a UNIX command-line program can be used to generate the file’s checksum (in Python the *hashlib*.*md5* command is used). For the validation of the FHR FASTA file with the checksum, FHR provides a command-line tool. An example of the command line tool is in Table 3.

**Table 3.**
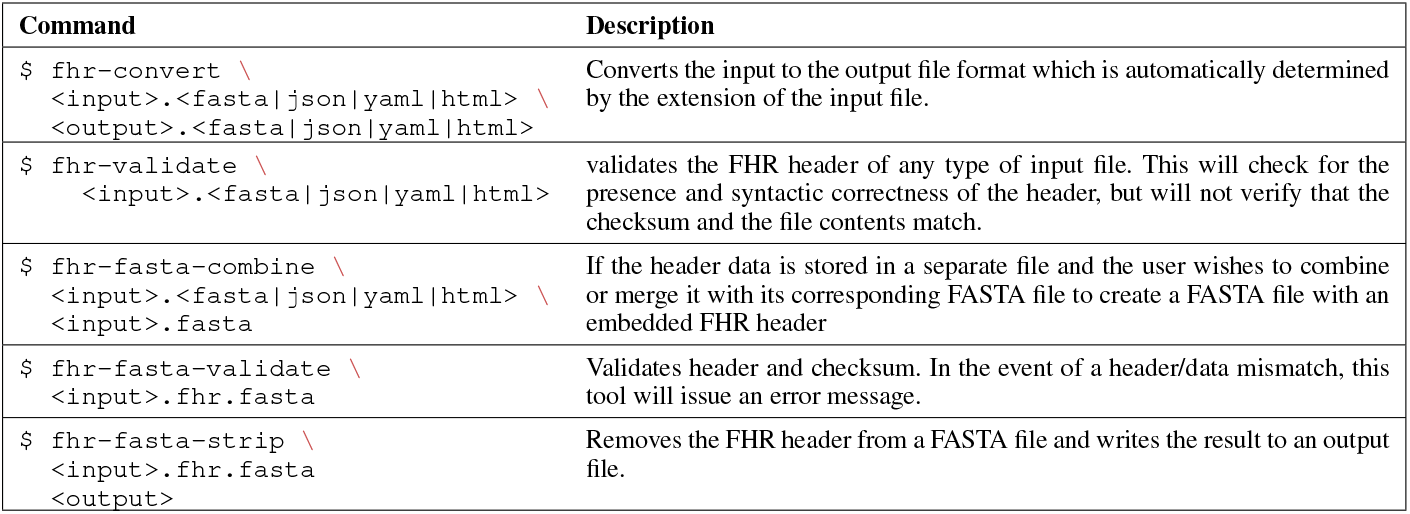
FHR Command Line.

**Table 4.** Software Locations for FHR Relevant Code.

| GitHub | DOI |
| --- | --- |
| FHR-Specification | 10.5281/zenodo.6762549 |
| FHR-File-Converter | 10.5281/zenodo.6762547 |
| Jbrowse (V1.7.9) with FHR support |  |

#### Software support

The schema code has been written to help developers work with FHR-specified reference genomes; the JSON validation tool is used to ensure that the FHR metadata conforms to the specification, a conversion tool to convert the FHR metadata into several different file formats, and the FHR software library used within applications written in Python.

### Conversion and validation validation tools

The FHR FASTA header can be validated using the JSON Schema (see: json-schema.org) using the FHR conversion and validation toolkit. This toolkit, which includes the FHR conversion and validation library, is written in Python. The Python library, in turn, depends on the Python libraries JSON, JSON-Schema, microdata, RE, and YAML. FHR tools provide several entry points into the Python library using the command line tools *fhr-convert* and *fhr-validate* and others (see Table 3). *fhr-convert* is for conversion between supported file formats, while *fhr-validate* is for validation of FHR metadata. The formats of the input and output files of the convert tool are automatically determined by the extension of the file. If the header data is stored in a separate file and the user wishes to combine or merge it with its corresponding FASTA file to create a FASTA file with an embedded FHR header, the user can use *fhr-combine-fasta*. To validate the FHR header of a file, use *fhr-validate*. This will check for the presence and syntactic correctness of the header but will not verify that the checksum and the file contents match. To validate the checksum also in an FHR FASTA file, the user would use *fhr-fasta-validate*. In the case of a header/data mismatch, this tool will issue an error message. If the user would like to remove the FHR header from a FASTA file, the *fhr-fasta strip* can be used. Examples of command line tools can be found in Table 3.

## DISCUSSION

FHR is designed to facilitate the transfer of provenance and metadata from external sources to or near genome assembly in a way that is minimally disruptive to existing bioinformatics workflows. To meet its goals of enabling the FAIR principles and reducing the risk of genome data and metadata divergence, FHR uses FASTA comments, keeps the included metadata to a minimum, provides a Python processing library, and includes alternative data serialisation. Additionally, FHR is written in Python, relies on JSON-schema for schema validation, and has a small codebase with few dependencies, making it less likely to enter an end-of-life stage in the near or medium future and reducing the maintenance burden. The only core dependency is the JSON schema, which itself relies on a JSON file that describes the schema rules, which is relatively small and can be easily converted if necessary.

### Achieving The Design Goals of the FHR format

FHR was designed with ease of implementation and backward compatibility in mind. Despite being described as “Minimal Information” models, many other metadata standards require a large number of fields and relationships, which makes implementation a challenge (see Taylor, 2007 (37) and Brazma, 2001 (38) for examples). Although these comprehensive standards theoretically offer robust inference capabilities, their level of adoption tends to be low due to the complexity of implementation, as information science theory might suggest (39). Following a design philosophy originating in the logic systems field (40, 41), FHR provides a fully featured metadata standard using as few fields as possible to achieve its goals. In an informal assessment of the time and effort needed to generate an FHR (Fragment Header) from the beginning, it was observed that an individual lacking familiarity with an assembly could proficiently produce a comprehensive FHR header within approximately 15 minutes.

Beyond ease of implementation, the FHR specification has four main design objectives:

1. Provide the user with the necessary metadata needed to replicate an analysis performed with a reference genome;
2. Support a variety of text representations;
3. Keep the metadata close to the data; and
4. Enable FAIR and TRUST. Furthermore, to ensure that data standards are beneficial to scientists and other stakeholders, it is necessary to design them to be user-friendly; and compatible with other standards and tools (42). Furthermore, ease of use and compatibility should be proportional to the utility of the new tool(43, 44, 45). Therefore, FHR is designed to fit into existing data and tool ecosystems to be adopted, and because it is located along the reference genome itself, it works more easily in a decentralised data environment.

#### Provide the metadata necessary to unambiguously identify the provenance of a genome

The FHR specification has been designed to provide the user with the data they need to trace the origin of a genome. This is done in two ways: by using a checksum to ensure the genome remains unchanged as it moves from one platform or device to another and by providing provenance metadata which can be used to reconstruct the continuity of the reference genome. For instance, the `genomeSynonym` field in FHR can be used to identify other names for the genome, or to link to related scholarly articles, authors, and instruments. Additionally, FHR provides information in the header to inform the user of the file’s contents, as well as links to external resources such as the NCBI.

#### Keep the metadata and data close

FHR ensures that metadata are easily accessible and correctly linked to the data by keeping data and metadata in the same file. The checksum in the FHR header is calculated on the basis of both the data and metadata. This allows the user to accurately identify the metadata and the data used in the analysis when using the checksum to identify the reference genome. This strong connection guarantees the origin of both the metadata and the data used in the analysis.

#### Support a variety of implementations

The principles of FHR design recognise that there are times when it is preferable to maintain the original FASTA file as is. Therefore, it is essential to offer a range of alternative serialisation techniques so that metadata can be stored alongside the data in the FASTA file. This is also beneficial in certain specialised circumstances, such as search engine optimisation, when we may want the metadata to be visible to a search engine without making the sequence data visible.

#### Enable FAIR and TRUST

FHR has been designed to ensure that those following the specification meet the core FAIR and TRUST principles. We designed the format with two user groups in mind. The first is the repository staff that provides the reference genomes to the community. The second group consists of researchers who use the reference genomes within their analytical pipelines. By supporting the FHR specification, the data repositories make the metadata more accessible to those using the files and will benefit from the users being able to find the data in the originating repository more readily. At the same time, the analytic pipelines written by researchers will become FAIRer and more TRUSTworthy by enabling the unambiguous identification of the genome(s) in use.

### Adoption Challenges

There are many software packages for manipulating FASTA files, but none currently understand FHR headers. Of more concern is the fact that many FASTA libraries do not follow the original standard’s use of the semicolon character to mark comment lines. To address this, the FHR toolset provides a command to strip the FASTA file of the FHR header when using software that cannot handle the header (see Table 3). In the future, we will work with the bioinformatics community to adopt standard pipelines to handle FHR-containing FASTA files. This will involve adding logic to existing FASTA software libraries to handle comments.

An unresolved issue is who has the authority to generate the FHR headers, which is a matter that needs to be debated by the stakeholder research communities. To minimise areas of authorship controversy, in the FHR header, the authorship of the assembly is separated from the authorship of the metadata, allowing a trusted third-party group, such as the curators of a model organism resource, to provide the metadata header.

#### Time and effort needed to create an FHR header

The time and effort required to develop an FHR header for a typical assembly is an important metric in the ease of adoption. To explore this question, we ran an example workflow in which we selected an unfamiliar mammalian genome assembly from an NCBI reference genome page (mouse GRCm39, RefSeq accession `GCF 000001635.27`), downloaded it, and created an appropriate FHR header using the data available on its NCBI genome page. In testing, the process of writing a single FHR header took 15 minutes, developing a template header JSON file to hold the information provided by NCBI. With automation, we estimate that the time per imported genome would be roughly five minutes.

#### Related Efforts

FHR is not the only project that attempts to solve the problem of aligning the storage of assembly metadata and the exchange of said data between resources. For example, the Minimum Information about a Genome Sequence (MIGS) specification, published in 2008, provides a set of fields for various types of assemblies with the intent of generating reports that are used to exchange information between resources (46). Compared to FHR, MIGS specification provides minimal information that should be tracked for a reference genome, whereas FHR only provides the fields that should be a header. Future versions of FHR may provide an extension mechanism that allows for the addition of fields from the full MIGS checklist.

FairGenomes is another project that tries to solve problems with metadata in Genomes (47). FairGenomes is designed for personal human genomes used in medical studies and takes advantage of a more extensive schema designed around using the stored metadata for personal human genomes in downstream analysis. FHR and FairGenomes have different design goals and use cases.

### Fostering community adoption

Data standards facilitate the rapid spread of concepts and ideas and become more useful the more researchers utilise them (ie, the network effect). To achieve widespread adoption, FHR needs early adopters to start generating and providing FHR-compliant reference genomes, while, in parallel, working on increasing the number of tools that can take advantage of additional information. Although it is still in its early days, the FHR header format has already been picked up and used by several significant biological curation efforts.

#### USDA ARS Pecan Germplasm phenotype database

The United States Department of Agriculture, Agricultural Research Service, Crop Germplasm Research Unit runs a Pecan Breeding program for pecan farmers, which contains phenotype measurements of living pecan trees and a library of nuts in its collection. The pecan phenotype database is being extended with genotype information that includes reference genomes of the pecan trees, and the metadata for the reference genomes will be determined by and exportable into FHR. In addition to the soft rollout of the Pecan phenotype and genotype database, other USDA projects are also starting to adopt FHR.

#### Alliance of Genome Resources

The Alliance of Genome Resources(29), a consortium of six Model Organisms and the Gene Ontology Consortium, is currently creating FHR YAML files. FHR YAML files are displayed within their JBrowse (48) genome browser instances.

#### AgBioData

AgBioData is a collection of Biological Resources with a mission to consolidate standards and is funded by the United States National Science Foundation. They are tasked with generating recommendations by acquiring, displaying, and reusing genomic, genetic, and breeding (GGB) data (49). AgBioData FAIR Scientific Literature and Genome Assembly and Annotation Nomenclature working groups are specifically orientated toward the problems of genome data and the metadata divergence problem.

We are working with this consortium as part of the FAIR Literature Working Group and the Genome Assembly Working Group to adopt FHR headers to use with the large number of genomes generated in agriculture-related projects.

#### MicroPublications

microPublication Biology is a peer-reviewed journal that revolutionizes the scholarly communication workflow by integrating data validation and curation into the publishing process. This curatorial-driven approach allows editors to review and authenticate domain-specific naming conventions and experimental reporting standards before publication. By working with the microPublication editorial team during publication, we will guide authors in properly reporting genome metadata standards by adopting FHR headers, alleviating numerous obstacles to making these data reusable.

#### KBase

KBase The Department of Energy Systems Biology Knowledgebase is currently testing its pipelines to see how well they handle files with FHR headers.

#### ATCC

ATCC’s Sequencing & Bioinformatics Center is testing the production of FHR compliant genome assemblies for the ATCC Genome Portal. Implementation into their production pipelines and inclusion in data deliverables to end-users is expected to be completed in 2024.

The goal of FHR is to provide a standard for maintaining the provenance of reference genomes that are translatable between storage and analysis platforms. To this end, FHR provides a schema of the minimal set of fields to facilitate the application of the FAIR and TRUST principles when generating and sharing Reference Genome files, as well as conversion and validation tools designed to make the creation, interconversion, and validation of FHR files easier.

## CONCLUSION

A metadata standard for reference genomes does not currently exist, and FHR represents a lightweight solution to this problem. FHR provides reference genomes with unambiguous identifiers, a method for validating the contents of the genome, and provenance identifying information, without placing an undue burden on the authors and maintainers of the genome. By providing several alternative implementations, FHR is minimally disruptive while providing many long-term benefits to the bioinformatics community. However, a transition to FHR will require the commitment of time and effort of various genomic data repositories and tool developers before its benefits will be realised by the bioinformatics community.

## DATA AVAILABILITY

The code published for FHR is in the public domain per the United States 17 U.S.C. § 105. The code and specification are freely available for use and modification (Table 4).

## ACKNOWLEDGEMENTS

The authors thank Natalie Meyers with The Lucy Family Institute for Data and Society at the University of Notre Dame for conversations around research communities focused on metadata standards that were used in the writing of the manuscript. The authors thank Monica Poelchau with the National Agriculture Library and Sarah Dyer at EMBL EBI for relevant discussions.

## FUNDING

Adam Wright is supported by the Adaptive Oncology Programme at the Ontario Institute for Cancer Research.

During a portion of this project, David Molik is supported by the USDA Agricultural Research Service (ARS) HQ Research Associate program in Big Data.

A portion of this work was carried out by the Tropical Pest Genetics and Molecular Biology Research Unit, ARS Project number 2040-22430-028-000D.

A portion of this work was carried out by the Arthropod-borne Animal Diseases Research Unit, ARS Project numbers 3020-32000-018-000-D, 3020-32000-020-000-D, and 3020-32000-019-000-D.

This research used resources provided by the SCINet project of the USDA Agricultural Research Service, ARS project number 0500-00093-001-00-D.

We gratefully acknowledge the support of the WormBase grant (U24HG002223), which provided funding for this research. This grant has been instrumental in supporting the contributions of Karen Yook, Daniela Raciti, Paul Sternberg, Adam Wright, Lincoln Stein, and Scott Cain. Their efforts have significantly contributed to the success of this project.

## AUTHORS NOTES

The U.S. Department of Agriculture is an equal opportunity lender, provider, and employer.

Mention of trade names or commercial products in this report is solely to provide specific information and does not imply recommendation or endorsement by the U.S. Department of Agriculture.

