## Supplemental Text for "Data Resources and Analyses Fair Header Reference genome: A Trustworthy standard"

### Supplement to FAIR Header Reference genome: A TRUSTworthy specification

*November 29, 2023*

**Abstract**

*This is the supplement to FAIR Header Reference genome: A TRUSTworthy specification*

#### 1 Supplement

##### 1.1 Broader Context behind FHR

In some way, when a data portal publishes data, it becomes the authority for the data it is publishing and takes on any responsibilities that the provision of those data to the larger scientific community entails [1, 2]. A decentralized data ecosystem is the disparate portals, communities, and resources that make up the emergent system a community member might use [3]. In the genomics space this might include NCBI resources [4], other data portals [5], conda environments [6], docker containers [7] and high-performance computing clusters. In a decentralised data ecosystem, it is up to the community to ensure that any data portal meets the minimum requirements for the community. Generally, this with the concept of trust [8], an example of trust in a data ecosystem would be if a bioinformatician uses a data resource online, they are implicitly trusting that the data resource is accurate or of sufficient quality. Another example of trust is when a data portal links to data in another portal which shows implicit trust in the other data portal. "trust" should be distinguished from TRUST principles, which are guidelines on how to build trust in digital repositories [9]. The more trust a data portal has, the more authority it has over that data [8]. One way to ensure that data is of sufficient quality is to record its provenance. Knowing the generation process behind a dataset helps to determine if the data is correct [8]. FHR provides a way to record provenance in such a decentralised ecosystem.

##### 1.2 FHR Implementation Notes

FHR has added the ability to provide a metadata file for the FASTA reference sequences to JBrowse, available as of release v1.7.9 (June, 2$^{\text{nd}}$ 2022). For the user to access this information, they need to click on "About Track" on the "Reference Track", after which a pop-up will appear displaying the header information. Although users can provide any text file for display, using the FHR specification, the user will more likely find the metadata more legible. In particular, the user may recognise the data being in the FHR format; not to mention that the file itself will more likely contain the appropriate information.

##### 1.2.1 FASTA Header Comments

Serialising with FASTA Comments merely involves reformatting the YAML formatted FHR metadata by adding a ";∼" prefix to each line and placing the resulting text at the top of the FASTA file. This serialisation has the added benefit of being directly associated with the reference genome. Although not commonly used, the original FASTA file specification allows for comments on lines prefixed by a semicolon. Many modern FASTA reading tools support the use of semicolon-denoted information in "legacy" mode. FHR utilises these FASTA comments to associate its metadata fields with the sequence itself; FHR tooling for FASTA formats looks for semicolon-prefixed lines in FASTA files, whereas most tools should based on the specification ignore these lines. FHR nevertheless recognises that such comments may exist in FASTA outside of their use to carry FHR-specified metadata. Therefore, to distinguish FHR metadata lines from other kinds of comments, FHR headers use a second character, the tilde, that is not part of the YAML or FASTA standards and therefore does not interfere with parsing these formats. The addition of the tilde should significantly reduce the chance of collisions in FASTA tooling and scripts which utilise regular expressions. Unfortunately, not all FASTA reading software can parse semicolon comments, even in legacy mode.

#### 1.3 FAIR

FAIR [10] and TRUST [9] were the guiding principles for the design of the FHR. FAIR and TRUST have checklists used to determine if a project meets their respective Guidelines (FAIR Checklist as part of Herczog et al. 2020 [11], for FAIR Principles see Table 1, modified from Wilkinson et al. 2016 [10], for TRUST Principles see Table 2, modified from Lin et al. 2020 [9]).

The FAIR principles were published in 2016 to improve the Findability, Accessibility, Interoperability, and Reuse (FAIR) of digital assets. These principles encourage the scholarly community to improve its usage and publication of metadata and intend to enhance the reuse of scientific datasets while focussing on machine actionability. Machine actionability is the amount to which a computational analysis can understand and make decisions on data, generally the combined concepts of machine readability, consistency, and the usefulness of the data to the analysis at hand. A focus on machine-actionability allows for automated data handling and validation. Metadata and unique persistent identifiers are fundamental to FAIR principles[10]. FHR has a full self-assessment of how it complies with the FAIR principles (see Table S.1).

#### 1.4 TRUST

In 2020, a complementary set of principles, "TRUST" for digital assets was published. This acronym stands for Transparency, Responsibility, User focus, Sustainability, and Technology. Although similar to the FAIR principles, these principles intend to provide a common framework to facilitate discussion and implementation of best practice in digital preservation by all stakeholders"[9]. FHR has a complete self-assessment of how it meets the TRUST principles (see Table S.2).

#### Table 1: Assessing FHR for FAIR Principles

| Principle | Criteria | Assessment |
|---|---|---|
| Findability | F1. (Meta)data are assigned a globally unique and persistent identifier<br>F2. Data are described with rich metadata (defined by R1 below)<br>F3. Metadata clearly and explicitly includes the identifier of the data they describe<br>F4. (Meta)data are registered or indexed in a searchable resource | • The field provided by FHR provides metadata to the user that goes beyond the genome name often located in the filename and links out to external resources providing further metadata<br>• FHR's checksum can be used as a globally unique ID<br>• FHR uses global CURIE'D identifiers to link to further information on the species and to external resources |
| Accessibility | A1. (Meta)data are retrievable by their identifier using a standardised communications protocol<br>A1.1 The protocol is open, free, and universally implementable<br>A1.2 The protocol allows for an authentication and authorisation procedure, where necessary<br>A2. Metadata are accessible, even when the data are no longer available | • The data is located directly inside the file with the genome itself. In this way, the information in the header is fully agnostic to the technology used to access it, including text editors.<br>• The file format does not inhibit data repositories from making the file freely available. format, item As a file format, the data provider can implement authorisation procedures to limit access to the data as appropriate. |
| Interoperability | I1. (Meta)data use a formal, accessible, shared, and broadly applicable language for knowledge representation.<br>I2. (Meta)data use vocabularies that follow FAIR principles<br>I3. (Meta)data include qualified references to other (meta)data | • The file FASTA headers themselves are in a non-binary text format which is well specified. The structure of the header follows YAML formatting making it both easy to read and computationally accessible.<br>• Tools are provided for converting between several file formats to interact with other systems<br>• The vocabulary chosen for the fields has been designed to align with fields from other technologies, with a focus on BioSchema.org fields. |
| Reuse | R1. (Meta)data are richly described with a plurality of accurate and relevant attributes<br>R1.1. (Meta)data are released with a clear and accessible data usage license<br>R1.2. (Meta)data are associated with detailed provenance<br>R1.3. (Meta)data meet domain-relevant community standards | • FHR specified files provide accurate and relevant attributes for describing the genome assembly.<br>• The checksum in the header can be used to uniquely identify the file for keeping provenance during analysis<br>• Currently the Bioinformatics community is using FASTA files without headers. Although adding the header may not be handled by some software the issue can be easily resolved through removing the header from the file before processing. |

#### Table 2: Assessing FHR for TRUST Principles

| Principle | Criteria | Assessment |
|---|---|---|
| Transparency | • Terms of use, both for the repository and the data holdings.<br>• Minimum digital preservation timeframe for the data holdings.<br>• Any pertinent additional features or services, for example, the capacity to responsibly steward sensitive data. | • The metadata is being stored with the data the user is less dependent on the data preservation timelines of data repositories<br>• The licence field allows the user to quickly determine the terms of use for the contents of the file<br>• By keeping a single file for both the data and metadata data is less likely to be mixed up, making it easier to make procedures for responsibly stewarding sensitive data |
| Responsibility | • Adhering to the designated community's metadata and curation standards, along with providing stewardship of the data holdings e.g. technical validation, documentation, quality control, authenticity protection, and long-term persistence.<br>• Providing data services e.g. portal and machine interfaces, data download or server-side processing.<br>• Managing the intellectual property rights of data producers, the protection of sensitive information resources, and the security of the system and its content. | • Genome repositories often house data in both databases and flat files. FHR-specified FASTA headers are intended to be handled by data repositories in a similar manner.<br>• Secondary files provide the opportunity to provide the FHR-specified metadata in commonly used file formats including HTML, JSON and YAML.<br>• File header information can be easily embedded in commonly used web interfaces including within JBrowse's reference track details.<br>• The licence field allows the intellectual property rights to be specified within the reference genome file. |
| User focus | • Implementing relevant data metrics and making these available to users.<br>• Providing (or contributing to) community catalogues to facilitate data discovery.<br>• Monitoring and identifying evolving community expectations and responding as required to meet these changing needs. | • FHR code is publicly hosted on GitHub. As the user community grows. In this way, the usage metrics will also be publicly available.<br>• The FHR tools and specifications are versioned and are intended to be updated as the needs of the community evolved |
| Sustainability | • Planning sufficiently for risk mitigation, business continuity, disaster recovery, and succession.<br>• Securing funding to enable ongoing usage and to maintain the desirable properties of the data resources that the repository has been entrusted with preserving and disseminating.<br>• Providing governance for necessary long-term preservation of data so that data resources remain discoverable, accessible, and usable in the future. | • Keeping the metadata in makes it so that no external technology is needed to access the data. In this way risks involving backing up and recovering the data is minimal<br>• As FHR has been designed to meet both FAIR and TRUST principles using the specifications should help satisfy requirements for securing funding<br>• As FHR is intended to evolve with the needs of the user community those using the technology will be able to implement the latest changes to continue to meet the needs of the community as well. |
| Technology | • Implementing relevant and appropriate standards, tools, and technologies for data management and curation.<br>• Having plans and mechanisms in place to prevent, detect, and respond to cyber or physical security threats. | • Tools are being provided to help implement the FHR standards including schema validation and serialization conversion tools.<br>• The FHR tools have minimal dependencies and are open source, minimising security risk. If security issues are detected within the dependencies, updates will be made to the code base to mitigate the issue. |

#### 1.5 Defining current Research Communities

There are many organisations tasked with maintaining reference genomes and related information. Two leading organisations mentioned earlier are the Alliance of Genome Resources and The i5k. Other notable projects are the Genome 10K Project [12], the Global Invertebrate Genome Alliance [13], and the 10,000 Plant Genomes Project [14].

##### 1.5.1 Defining Alliance of Genome Resources

The Alliance[15] provides a portal for curated and computationally derived information for many of the main model organisms. It is made up of the following founding members: FlyBase (FB), Gene Ontology (GO), Rat Genome Database (RGD), Mouse Genome Database (MGD), Saccharomyces Genome Database (SGD), WormBase (WB), and the Zebrafish Network (ZFIN).

The Alliance makes reference genomes available for download and viewing through JBrowse (see: www.alliancegenome.org/about-us). Providing a metadata file for download or through the JBrowse user interface would allow the users to access the provenance of the genome assembly they are observing in a standardised format.

##### 1.5.2 Defining i5k

i5k, Sequencing Five Thousand Arthropod Genomes project [16, 17]. One of the sizable members of the i5k project is the ag100pest initiative [18]. The i5k project has four main goals: to sequence and analyse 5000 arthropod species; to provide guidelines and best practices to other arthropod genome projects; to help existing and new arthropod projects find appropriate repositories; and to grow a community that works toward improving sequencing, assembly, annotation, and data management standards (see: i5k.github.io/about). The i5k also makes reference genomes available for download and viewing through JBrowse (Apollo derivative).

#### 1.6 JSON Schema in Full

#### 1.7 Alternative Data Serialisations

FHR has three alternative data serialisations that can be converted to and from, for specific use cases where a FASTA header may not be possible. Alternative formats can be found in FAIR-BioHeaders/FHR-Specification and FAIR-BioHeaders/FHR-File-Converter. FAIR-bioHeaders provides tools for converting the FASTA header information into several file formats: YAML, JSON, and Microdata. Each file format has specific use cases. with YAML FFRGS is human-readable, with microdata, embeddable, and with JSON for more interoperable. Rather than being a hindrance to the standardisation of reference genome provenance metadata, additional data serialisations provide unique benefits. Additionally, the format conversion tool increases interoperability. These supplementary files should have the same name as the reference genome FASTA file with file extension from <genome file name>.fasta to .fhr.<yaml|json|html> and should be located together with the FASTA file itself. It is important to co-locate these files with the FASTA file if it is impossible to place the header in the FASTA file itself, especially for FAIR compliance.

Figure 1: FHR Schema Version One in Full

```
{
  "$schema": "https://json-schema.org/draft/2020-12/schema",
  "$id": "https://raw.githubusercontent.com/FAIR-bioHeaders/FHR-
     Specification/main/fhr.json",
  "title": "FHR",
  "description": "FAIR Header Reference genome Schema for Genome
     Assemblies",
  "type": "object",
  "properties": {
    "schema":{
      "type": "string",
      "description": "centralized schema file"
    },
    "schemaVersion":{
      "type": "number",
      "description": "Version of FHR"
    },
    "genome":{
      "type": "string",
      "description": "Name of the Genome"
    },
    "genomeSynonym": {
      "type": "array",
      "description": "Other common names of the genome",
      "items": {
        "type": "string"
      }
    },
    "taxon":{
      "type": "object",
      "description": "Species name and URL of the species
         information at identifiers.org",
      "properties": {
        "name": {
          "type": "string"
        },
        "uri": {
          "type": "string",
          "format": "uri",
          "pattern": "https://identifiers.org/taxonomy:[0-9]+"
        }
      }
    },
    "version":{
      "type": "string",
      "description": "Version number of Genome eg. 1.2.0"
    },
    "metadataAuthor":{
      "type": "array",
      "items": {
        "type": "object",
        "description": "Author of the FFRGS Instance (Person or
           Org)",
        "properties": {
          "name": {
```

Figure 2: Example Minimal FHR YAML file


```yaml
schema: https://raw.githubusercontent.com/FAIR-bioHeaders/FHR-
   Specification/main/fhr.json
schemaVersion: 1.0
genome: Example species
taxon:
  name: Example species
  uri: https://identifiers.org/taxonomy:0000
version: 0.0.1
metadataAuthor:
  name: Adam Wright
  uri: https://orcid.org/0000-0002-5719-4024
assemblyAuthor:
  name: David Molik
  uri: https://orcid.org/0000-0003-3192-6538
dateCreated: '2022-03-21'
checksum: md5:7582b26fcb0a9775b87c38f836e97c42
```


##### 1.7.1 YAML file

YAML is a data format intended to be human-readable [19] and is therefore intended to be the primary FFRGS serialisation, either as a secondary file or as the preferred part of a FASTA header.

##### 1.7.2 JSON file

JSON is by far the most dominant data exchange format for interprocess communication on the web and is relatively typical as a standalone data storage format [20]. JSON started as a data exchange format for JavaScript [21] applications, a popular web scripting language that can produce interactive websites and visualisations. Over time, JSON has become ubiquitous across languages and technologies.

##### 1.7.3 Microdata

Microdata is a tagging implementation to put metadata into HTML code [22]. By embedding the FFRGS standardised metadata into the HTML of a Web page, the information is directly available to search engines and for others to scrape from the contents of the page.

Figure 3: Example Minimal FHR JSON file

```
{
    "$schema": "https://raw.githubusercontent.com/FAIR-bioHeaders
       /FHR-Specification/main/fhr.json",
    "$id": "https://raw.githubusercontent.com/FAIR-bioHeaders/FHR
       -Specification/main/examples/example.fhr.json",
    "schema": "https://raw.githubusercontent.com/FAIR-bioHeaders/
       FHR-Specification/main/fhr.jso",
    "schemaVersion": 1.0,
    "taxon": {
      "name": "Example species",
      "uri": "https://identifiers.org/taxonomy:0000"
    },
    "genome": "Example species",
    "version": "0.0.1",
    "metadataAuthor": [
      { "name": "Adam Wright",
      "uri": "https://orcid.org/0000-0002-5719-4024" }
    ],
    "assemblyAuthor": [
      { "name": "David Molik",
      "uri": "https://orcid.org/0000-0003-3192-6538" }
    ],
   "dateCreated": "2022-03-21",
    "checksum": "md5:7582b26fcb0a9775b87c38f836e97c42"
}
```

Figure 4: Example Minimal FHR Micodata

```
<div itemscope itemtype="https://raw.githubusercontent.com/FAIR-
   bioHeaders/FHR-Specification/main/fhr.json" version="1">
  <span itemprop="schema">https://raw.githubusercontent.com/FAIR-
     bioHeaders/FHR-Specification/main/fhr.json</span>
  <span itemprop="schemaVersion">1.0</span>
  <span itemprop="version">0.0.1</span>
  <span itemprop="genome">Example species</span>
  <span itemprop="metadataAuthor">
    <span itemprop="name">Adam Wright</span>
    <span itemprop="uri">https://orcid.org/0000-0002-5719-4024</
       span>
  </span>
  <span itemprop="assemblyAuthor">
    <span itemprop="name">David Molik</span>
    <span itemprop="uri">https://orcid.org/0000-0003-3192-6538</
       span>
  </span>
  <span itemprop="taxon">
    <span itemprop="name">Bomnas huntii</span>
    <span itemprop="uri">https://identifiers.org/taxonomy:9606</
       span>
  </span>
  <span itemprop="checksum">md5:7582b26fcb0a9775b87c38f836e97c42<
     /span>
</div>
```

#### Disclaimer

The U.S. Department of Agriculture is an equal opportunity lender, provider, and employer.

Mention of trade names or commercial products in this report is solely for the purpose of providing specific information and does not imply recommendation or endorsement by the U.S. Department of Agriculture.
